# Oxidative stress in retinal pigment epithelial cells increased endogenous complement-dependent inflammatory and angiogenic responses - independent from exogenous complement sources

**DOI:** 10.1101/722470

**Authors:** Timon-Orest Trakkides, Nicole Schäfer, Maria Reichenthaler, Konstanze Kühn, Volker Enzmann, Diana Pauly

**Author notes:** corresponding author, University Hospital Regensburg, Eye clinic, Experimental Ophthalmology, Franz-Josef-Strauss-Allee 11, Regensburg, 93053, Germany.

## Abstract

Oxidative stress-induced damage of the retinal pigment epithelium (RPE) together with chronic inflammation has been suggested as major contributors to retinal diseases. Here, we examine the effects of oxidative stress and endogenous complement components on the RPE and its pro-inflammatory and –angiogenic responses.

The RPE cell line, ARPE-19, treated with H_2_O_2_ reduced cell-cell contacts, increased marker for epithelial–mesenchymal transition but showed less cell death. Stressed ARPE-19 cells increased the expression of complement receptors CR3 and C5aR1. CR3 was co-localized with cell-derived complement protein C3, which was observed in its activated form in ARPE-19 cells. C3 as well as its regulators CFH and properdin accumulated in ARPE-19 cells after oxidative stress independent from external complement sources. This cell-associated complement accumulation promoted *nlrp3* and *foxp3* expression and subsequent increased secretion of pro-inflammatory and pro-angiogenic factors. The complement-associated ARPE-19 reaction to oxidative stress, independent from external complement source, was increased by the PARP-inhibitor olaparib.

Our results indicated that RPE cell-derived complement proteins and receptors are involved in RPE cell homeostasis following oxidative stress and should be considered as targets for treatment developments for retinal degeneration.

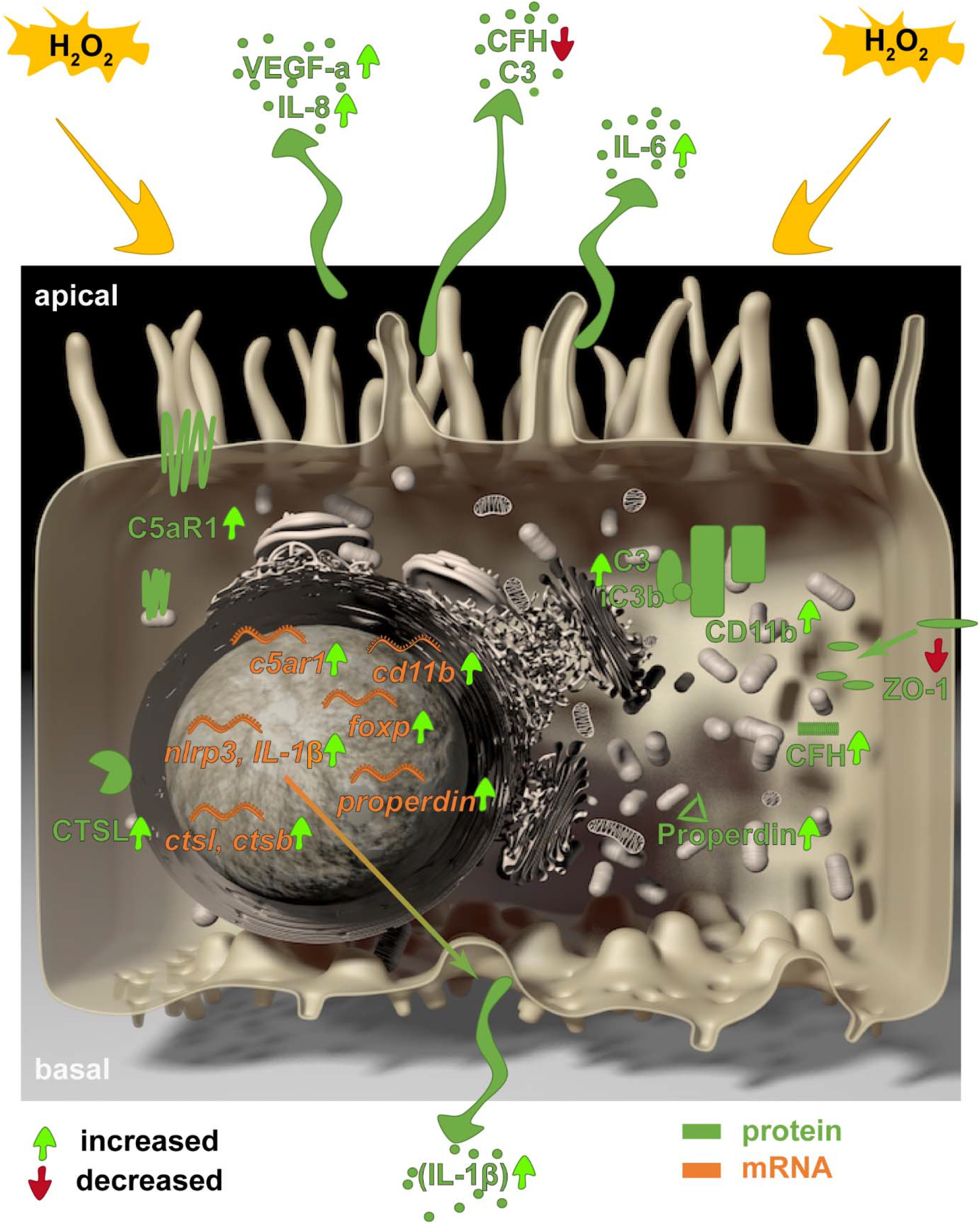
GRAPHICAL ABSTRACT.

We show a functional link between oxidative stress, complement receptors, endogenous complement proteins, pro-angiogenic and -inflammatory responses in ARPE-19 cells. These effects are independent from extracellularly added complement proteins or receptor ligands. We suggest an oxidative stress-associated autocrine mechanism of complement receptor regulation in ARPE-19 cells in connection with upregulated intracellular proteases.

**HIGHLIGHTS:**

- Oxidative stress accumulates complement proteins and receptors in RPE cells
- Oxidative stress activates the RPE inflammasome without external complement proteins
- Oxidative stress increases *foxp3* expression and IL-8/VEGF secretion in RPE cells
- Olaparib enhances pro-inflammatory response of RPE

## INTRODUCTION

One of the most oxidative environments in the body is the retinal pigment epithelium (RPE) [1], which is in close contact with the photoreceptors and maintains visual function [2]. Low levels of reactive oxygen species are required to maintain physiological functions [3], but the combination of exposure to visible light, elevated metabolic activity, accumulation of oxidized lipoproteins and decreased antioxidant functions during aging make the retinal tissue vulnerable to oxidative stress [4,5]. Oxidative damage to the RPE was therefore identified as a contributing factor to different retinal degenerative diseases such as age-related macular degeneration or Stargardt disease [6–8].

In line with this, chronic oxidative stress can involve chronic inflammation subsequently leading to cellular damage in the RPE/retina [6,9]. Based on genetic polymorphisms in genes of the complement system, systemic complement activation and local complement deposition in degenerative retinal tissue a contribution of the complement system to oxidative stress-related retinal degeneration was hypothesized [7,10,11]. The complement system is composed of over 40 proteins, which bridge the innate and adaptive immune defence [12]. The main functions are (I) removal of damaged cells, (II) protection against invading pathogens and (III) attraction of immune cells.

Beside the traditional view, evidence is accumulating that complement is also involved in physiological processes such response to oxidative stress and cellular survival programmes [6]. The complement system comprises several soluble and membrane-bound proteins and receptors, which can be produced by a number of cells, including non-immune cells and extrahepatic tissue, and contribute to the autocrine cell physiology [13]. The role of endogenous complement-dependent regulation of cellular homeostasis has been extensively studied in the recent years in T-cells [14]. T-cells, B-cells and human airway epithelial cells contain intracellular stores of C3, which is endogenously cleaved into its active forms C3a and C3b by intracellular proteases [15–17]. Activated C3 was correlated with the activation of the NLRP3 inflammasome in T-cells [15], which lead to chronic pro-inflammation. An antagonising complement modulation was described for regulatory T-cells, where C3aR and C5aR1 activation resulted in the activation of the forkhead box P3 (FOXP3) transcription factor [15,22]. The FOXP3 transcription factor acts in multimodal fashion and stimulates the release of anti-inflammatory cytokines and pro-angiogenic factors [22–24].

Oxidative stress and inflammasome activation were previously correlated to external complement activity in RPE cells [6,25]. FOXP3 activation in RPE cells also depended on extracellularly added complement components [26]. However, RPE-derived complement has not been discussed as source for NLRP3 or FOXP3 modulation. Complement components can be produced by RPE cells [27] and their expression is changed under oxidative stress [28–32]. Further activated forms of C3 (C3a), independent from extracellular complement sources, were also secreted by RPE cells, suggesting a similar function of the complement system in RPE cells compared to T-cells [18–21].

In this study, we report that H_2_O_2_ stimulated the accumulation of complement protein C3, CFH and properdin in RPE cells and increased the expression of complement receptors C5aR1 and CR3. This was accompanied with increased NLRP3 inflammasome activation and FOXP3-associated release of pro-angiogenic factors. Our results indicate a cell homeostatic function of cell-derived complement components, independent from external complement receptor ligands.

## RESULTS

### Stressed, *in vivo*-like cultivation of ARPE-19 cells

We investigated cellular stress response and cell-specific complement expression in a cell line of human RPE cells, the ARPE-19 cell line. Aged ARPE-19 cells of passage 39 were cultivated under *in vivo*-like, unstressed conditions. This was visualized by staining of *zonula occludens* 1 (ZO-1), an important protein for cell-cell-contact, showing formation of stable tight junctions and mainly mononuclear, polarized cell growth on transwell filters (Fig. 1A, D). H_2_O_2_ treatment resulted in cellular stress indicated by reduced cell-cell contacts after 4 h (Fig. 1B) and a time-dependent translocation of ZO-1 from the cell membrane to the cytoplasm after 24 h (Fig. 1E). Evidence of induced cellular stress by H_2_O_2_ were also observed by increased mRNA expression of vimentin (*vim*) and α smooth muscle actin (*α-sma*), typical mesenchymal marker indicating epithelial–mesenchymal transition (Sup. Fig. 1) [33–35]. However, the majority of the ARPE-19 cells did not undergo apoptosis under these non-lethal oxidative stress conditions, shown by a low number of TUNEL-positive cells (Fig. 1C, F).

**Fig. 1.**
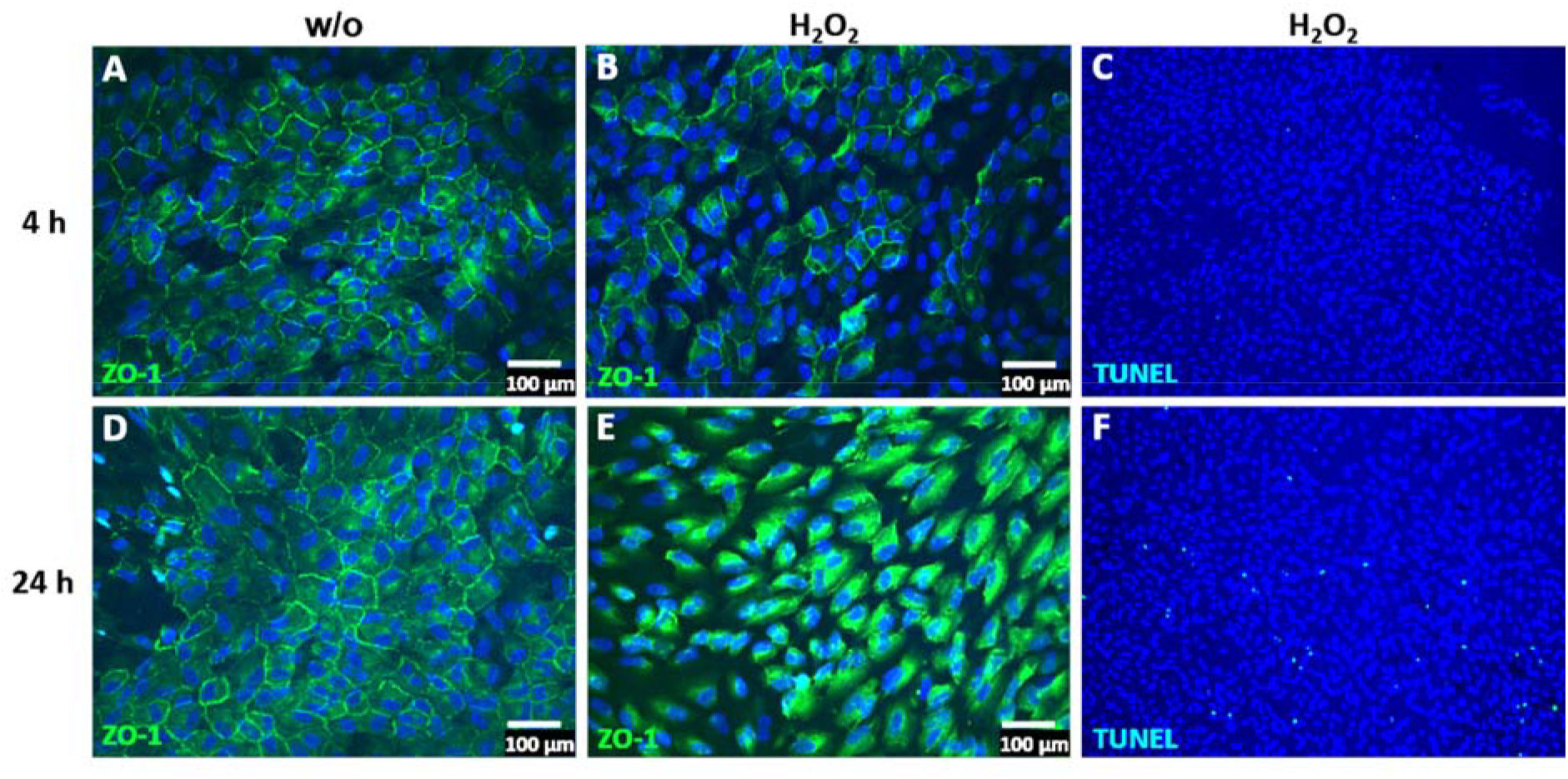
ARPE-19 cells reduced tight junctions and circumvent apoptosis under oxidative stress. **(A, D)** ARPE-19 cells untreated (w/o) and stressed with H_2_O_2_ for **(B, C)** 4 h or **(E, F)** 24 h translocated time-dependently the *zonula occludens* protein 1 (ZO-1, green) from the **(A, D)** cell membrane to the **(B, E)** cytoplasm. **(C, F)** ARPE-19 cells treated with oxidative stress showed a minimal TUNEL-positive (light blue) apoptotic reaction after **(F)** 24 h.

### ARPE-19 cells increase complement receptor expression under oxidative stress

ARPE-19 cells express cellular receptors, sense the cellular environment and can react to complement activation products. Complement receptor 3 (CR3), is a heterodimer integrin consisting of two non-covalently linked subunits CD11b and CD18 on leukocytes/ microglia and is activated by C3 cleavage products (iC3b, C3d, C3dg). CD11b has been detected with low expression on mRNA and protein level in ARPE-19 cells (Fig. 2A, B). Oxidative stress increased the *cd11b* mRNA expression after 4 h, which was also shown on protein level with immunostaining (Fig. 2A, C).

**Fig. 2.**
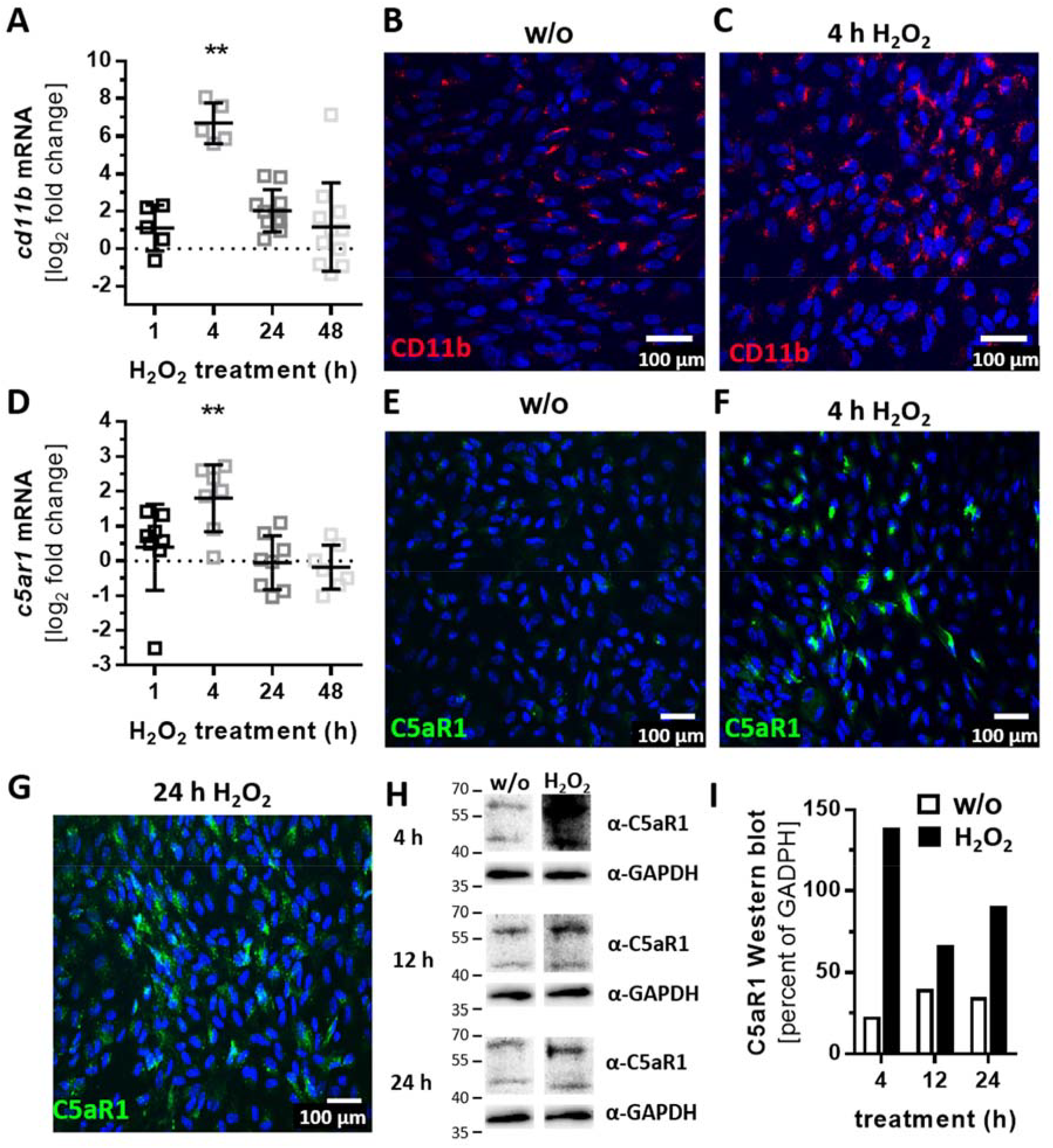
Oxidative stress increased expression of complement receptor subunit CD11b and C5aR1 in ARPE-19 cells. **(A, D)** *Cd11b* and *c5aR1* mRNA expression was increased 4 h following H_2_O_2_ treatment. This effect was confirmed on protein level by immunohistochemistry using **(B, C)** anti-CD11b (red) and **(E – G)** anti-C5aR1 (green) antibodies. **(H)** Western Blots of ARPE-19 cell lysates detected C5aR1 between 40 – 60 kDa after 4 – 24 h H_2_O_2_ treatment. **(I)** Quantitatively, C5aR1 expression was increased in H_2_O_2_ treated cells in Western blots. **(A, D)** Mean with standard deviation is shown, ** p≤ 0.01 unpaired, two-tailed, parametric t-test, dotted line depicts untreated control, **(B, E, H, I)** w/o untreated control.

Activation of complement protein C5 is detected by complement receptor C5aR1, which is expressed by ARPE-19 cells (Fig. 2D). H_2_O_2_ treatment increased *c5ar1* expression comparable to *cd11b* expression (Fig. 2D – F). C5aR1 protein accumulation was observed after 4 h at the cell nuclei (Fig. 2F), which was more distributed in/on the cell after 24 h (Fig. 2G). Increased C5aR1 protein level was also confirmed in Western blots (Fig. 2H, I).

Transcription levels of complement receptor *c3aR* was not significantly changed in H_2_O_2_-treated ARPE-19 cells (Sup. Fig. 2A).

### Complement proteins accumulated in ARPE-19 cells under oxidative stress

Complement proteins, which can modulate the activity of complement receptors at the RPE, are locally produced in the retina [27,36] and by RPE cells (Fig. 3, Sup. Fig. 2B – K). The mRNA expression and protein levels of the stabilizing complement regulator properdin were increased after 24 h of H_2_O_2_ treatment (Fig. 3A – D), but apical properdin secretion was not detected (Sup. Fig. 3A). This indicated a properdin storage in the stressed ARPE-19 cells (Fig. 3B – D).

**Fig. 3.**
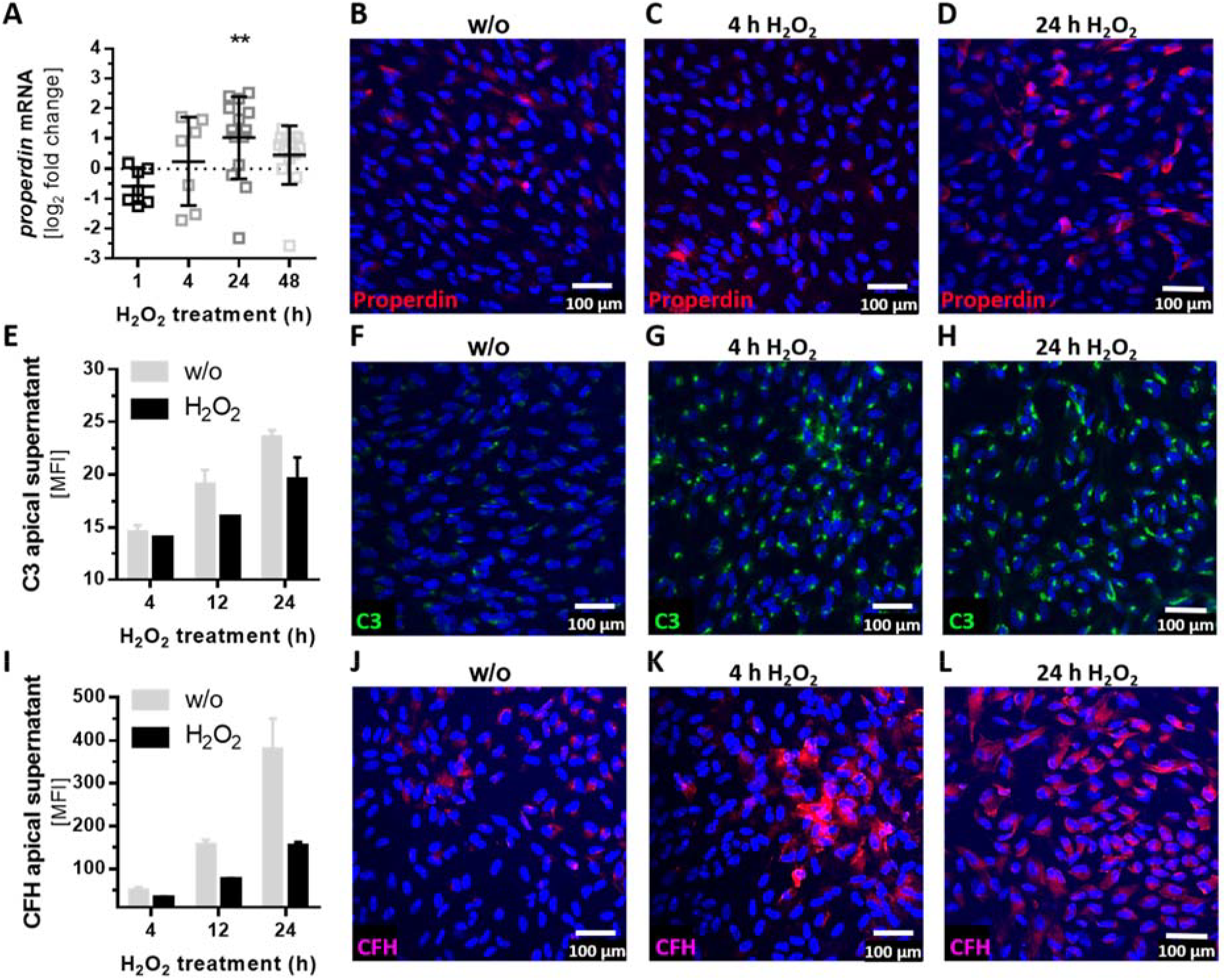
Oxidative stress induced complement component accumulation in ARPE-19 cells. **(A)** The *properdin* mRNA level was increased 24 h following H_2_O_2_ treatment. This effect was confirmed on protein level by immunohistochemistry using an **(B – D)** anti-properdin (red) antibody. **(E)** C3 and **(I)** CFH protein concentration in the apical supernatant of ARPE-19 cells decreases following H_2_O_2_ treatment. Immunohistochemistry using **(F – H)** anti-C3 (green) and **(J – L)** anti-CFH (purple) antibodies showed an increase of cell-associated **(G, H)** C3 and **(K, L)** CFH after oxidative stress treatment. **(A, E, I)** Mean with standard deviation is shown, **(A)** ** p≤ 0.01 unpaired, two-tailed, parametric t-test, dotted line depicts untreated control, **(E, B, F, I, J)** w/o untreated control.

Transcription levels of additionally tested complement components (*c3*, *c4a*, *c4b*, *cfb*, *cfd*, *c5*), soluble (*cfh*, *cfi*) and membrane-bound complement regulators (*cd46*, *cd59*) did not change under oxidative stress conditions (Sup. Fig. 2B – K).

However, we observed a change in cellular accumulation and modulated secretion of complement components on protein level by oxidative stress (Fig. 3E – L). Central complement component *c3* was not regulated on mRNA expression by oxidative stress (Sup. Fig. 2B), but we detected an increase of cellular C3 in immunostainings and a decrease of C3 secretion into the apical supernatant of ARPE-19 cells (Fig. 3E – H). Secretion of C3 was more observable in younger compared to older ARPE-19 cells treated with H_2_O_2_ (Sup. Fig. 3B). A similar effect of cellular complement component accumulation and reduced secretion was detectable for complement regulator CFH (Fig. 3I – L, Sup. Fig. 3C). Thus, the mRNA expression was not changed under oxidative stress (Sup. Fig. 2H).

### Autocrine complement receptor activation following oxidative stress is correlated with the release of pro-inflammatory and pro-angiogenic factors

Intracellular complement proteins and cellular complement receptors were previously associated with an autocrine regulation of cell differentiation and cell physiology in T-cells as well as lung epithelial cells [17,37]. In line with this we found a co-localization of CD11b and C3 in ARPE-19 cells (Fig. 4A, B) and activated C3 fragments (C3b α’, C3d) in the ARPE-19 cells (Fig. 4C), without adding any external complement source.

**Fig. 4.**
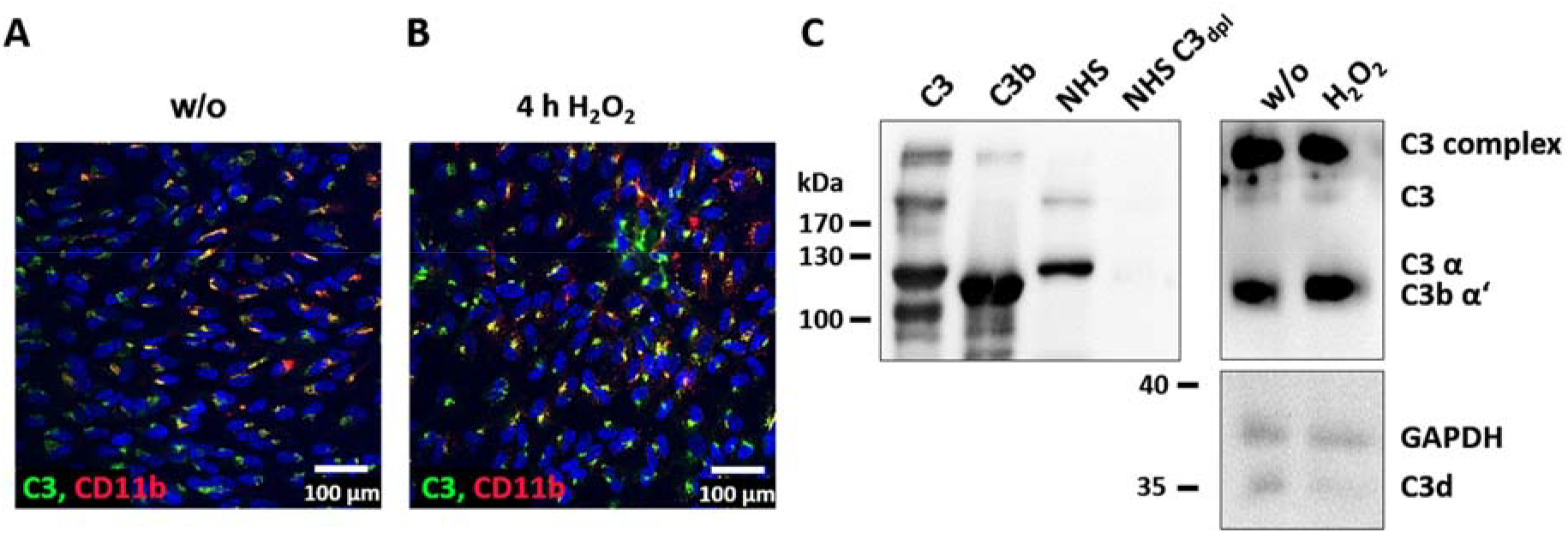
C3 and complement receptor CR3 are co-localized in ARPE-19. **(A)** Unstressed (w/o) and **(B)** H_2_O_2_ treated ARPE-19 cells were stained with anti-C3 (green) and anti-CD11b (red) antibodies. Overlapping staining signals (yellow) suggested a co-localization of C3 and CD11b. **(C)** C3 and activation products (C3b α’ and C3d), were detected in untreated and H_2_O_2_ treated ARPE-19 cells, together with native C3, C3b, human serum (NHS) and C3 depleted human serum (NHS C3_dpl_), using Western Blot under reducing conditions.

Intracellular cleavage of complement proteins into active fragments, independent from the systemic complement cascade, can be mediated by intracellular proteases, as cathepsin B (CTSB) or cathepsin L (CTSL) [14,15]. Both proteases were expressed by ARPE-19 cells and they were upregulated following oxidative stress (Fig. 5). The expression of *ctsb* and *ctsl* mRNA was increased after 24 h of H_2_O_2_ treatment (Fig. 5A, B). We confirmed the higher concentration of CTSL in ARPE-19 cells under stress conditions also on protein level (Fig. 5C, D).

**Fig. 5.**
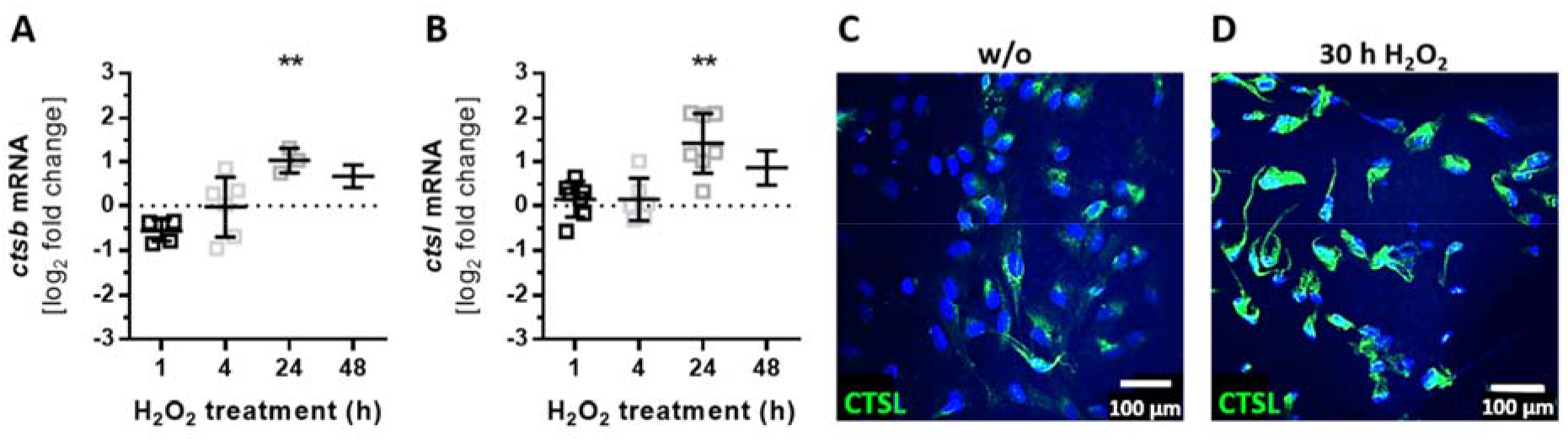
Expression of intracellular proteases is increased by oxidative stress in ARPE-19. **(A)** *Ctsb* and **(B)** c*tsl* mRNA expression increased 24 h following H_2_O_2_ treatment. This effect was confirmed on protein level in immunostainings using an **(C, D)** anti-CTSL (green) antibody. **(A, B)** Mean with standard deviation is shown, ** p≤ 0.01 unpaired, two-tailed, parametric t-test, dotted line depicts untreated control, **(C)** w/o untreated control.

Activation of complement receptors on the one hand can induce inflammasome activation and on the other hand can regulate the mTOR-pathway involving the FOXP3 transcription factor in T- and RPE cells [25,26,38], the well-coordinated interplay of complement receptor signalling controls the pro- and anti-inflammatory cytokine release [25,39]. After detection of cell-derived C3 co-localized with CD11b, its activation products C3b and C3d (Fig. 4) as well as H_2_O_2_ dependent regulation of complement receptors (Fig. 2) and cellular complement protein accumulation (Fig. 3), we supposed also an autocrine, complement-dependent role of the NLRP3 inflammasome and FOXP3 in ARPE-19 cells treated with H_2_O_2_ to induce oxidative stress. This regulation would be independent of blood-derived complement components and involves release of cytokines and growth factors in stressed ARPE-19 cells (Fig. 6).

**Fig. 6.**
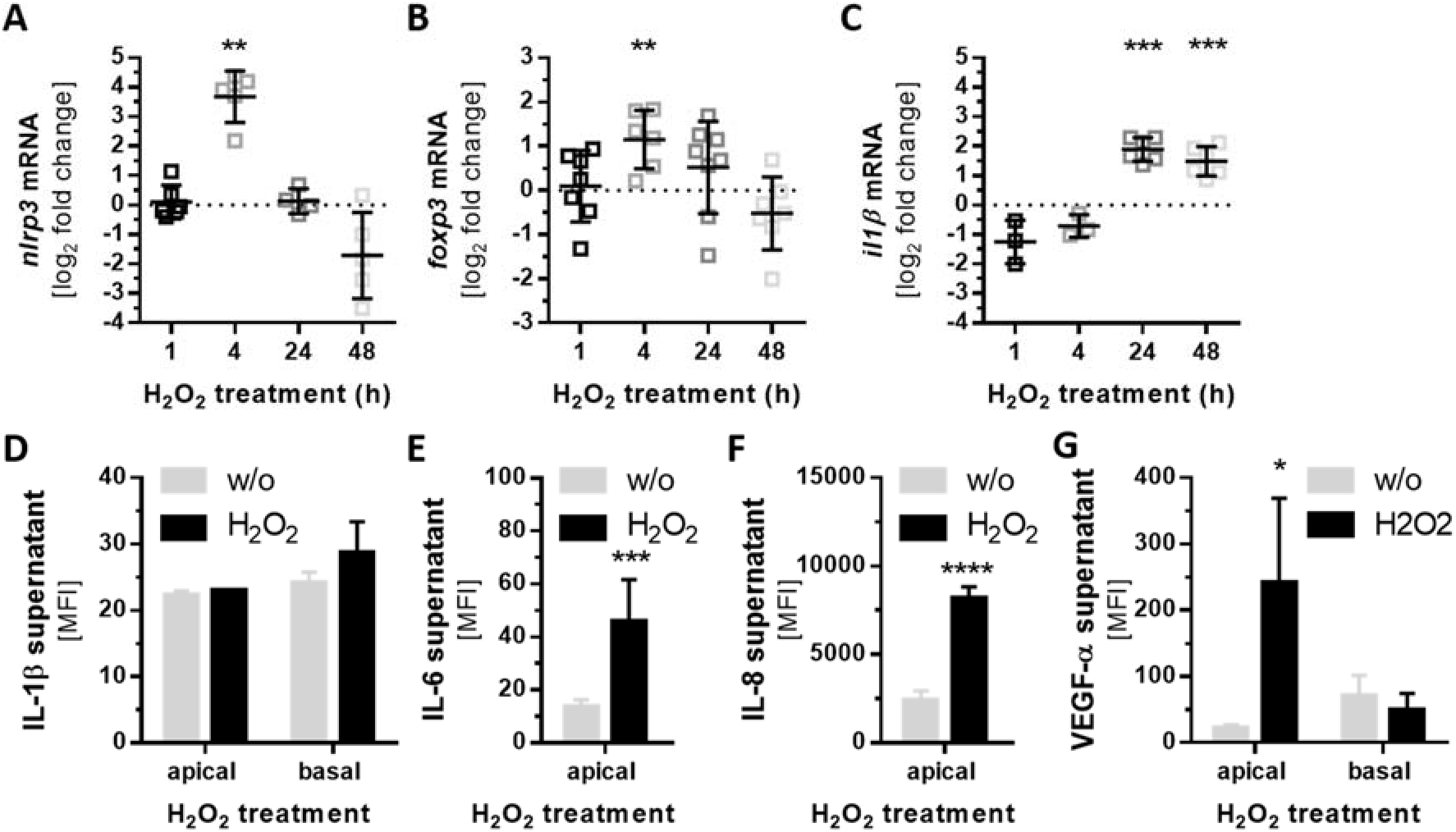
Increased *nlrp3* and *foxp3* mRNA expression correlates with pro-inflammatory and pro-angiogenic factor secretion. **(A)** *Nlrp3*, **(B)** *foxp3* and **(C)** *il1β* mRNA levels increased either **(A, B)** 4 h or **(C)** 24 h and 48 h following H_2_O_2_ treatment. Pro-inflammatory cytokine release of **(D)** IL-1β (after 4 h) and **(E)** IL-6 (after 48 h) was detected in stressed ARPE-19 cells. This was correlated with an enhanced secretion of pro-angiogenic factors **(F)** IL-8 (after 48 h) and **(G)** VEGF-α (after 4 h) H_2_O_2_ treated cells. MFI mean fluorescence intensity. Mean with standard deviation is shown, * p≤ 0.05, ** p≤ 0.01, *** p≤ 0.001, **** p≤ 0.0001 unpaired, two-tailed, parametric t-test, **(A, B, C)** dotted line depicts untreated control, **(D ̶ G)** w/o untreated control.

Indeed, we detected an increased expression of *nlrp3* and *foxp3* mRNA after 4 h of H_2_O_2_ treatment (Fig. 6A, B). A subsequent enhanced expression of *il1β* mRNA after 24 h and 48 h indicated the activation of the NLRP3-inflammasome in stressed ARPE-19 cells (Fig. 6C), thus the mRNA expression of *il18* was not changed (Sup. Fig. 2L). Consequently, we found higher pro-inflammatory cytokine levels in H_2_O_2_ treated ARPE-19 cell supernatant compared to untreated control (Fig. 6D – E). IL-1β was slightly increased shortly after treatment (4 h), while IL-6 was significantly elevated in supernatant of stressed RPE cells (Fig. 6D).

Increased *foxp3* expression is an attribute of anti-inflammatory regulatory T-cells, which secrete mainly TGF-β and IL-10. We did not detect a change in *tgfβ* expression (Sup. Fig. 2M) or IL-10 secretion (data not shown) by H_2_O_2_ treated ARPE-19 cells. Therefore, we assumed a pro-angiogenic function of *foxp3* in the cells as previously reported [23,24]. In line with this, we observed an increase of IL-8 (after 48 h) and VEGF-α (after 4 h) secretion in stressed ARPE-19 cells (Fig. 6F, G). This correlation between complement components, *foxp3* expression and pro-angiogenic reaction in RPE cells needs to be further investigated.

[As a side note: IL-17, IFN γ, IL-18, IL-2 and TNF-α were not detected in the apical and basal supernatant of 4 h, 24 h and 48 h untreated and H_2_O_2_treated ARPE-19 cells (data not shown).]

### Olaparib boosted the pro-inflammatory response of ARPE-19 cells to oxidative stimuli

Oxidative stress-induced cellular reactions were previously ameliorated by an approved anti-cancer drug olaparib, which is an inhibitor of the poly(ADP-ribose) polymerase (PARP) [40–42]. We investigated the effect of olaparib on H_2_O_2_-dependent mRNA expression changes of complement receptors, components and inflammation-related transcripts (Fig. 7, Sup. Fig. 4). Oxidative stress increased expression of *cd11b*, *c5ar1* and *nlrp3* after 4 h of H_2_O_2_ treatment, this was further enhanced by olaparib-treatment (Fig. 7A, B, C). An increase of *properdin* and *ctsb* transcripts was observed after 24 h following oxidative stress alone (Fig. 3A, 5A). A combination of H_2_O_2_ and olaparib accelerated this reaction with a significant increase of *properdin* and *ctsb* mRNA expression already after 4 h (Fig. 7D, E). The expression of *cfd* (Sup. Fig. 2F) was not modulated under oxidative stress, however H_2_O_2_ and olaparib together increased the *cfd* transcript level (Fig. 7F). Olaparib did not interfere with transcription of *foxp3* (Fig. 7G) and other transcripts (*c3*, *c4a*, *c5*, *cfb*, *cfh*, *cfi*, *c3ar*, *ctsl*) (Sup. Fig. 4) in ARPE-19 cells treated with H_2_O_2_.

**Fig. 7.**
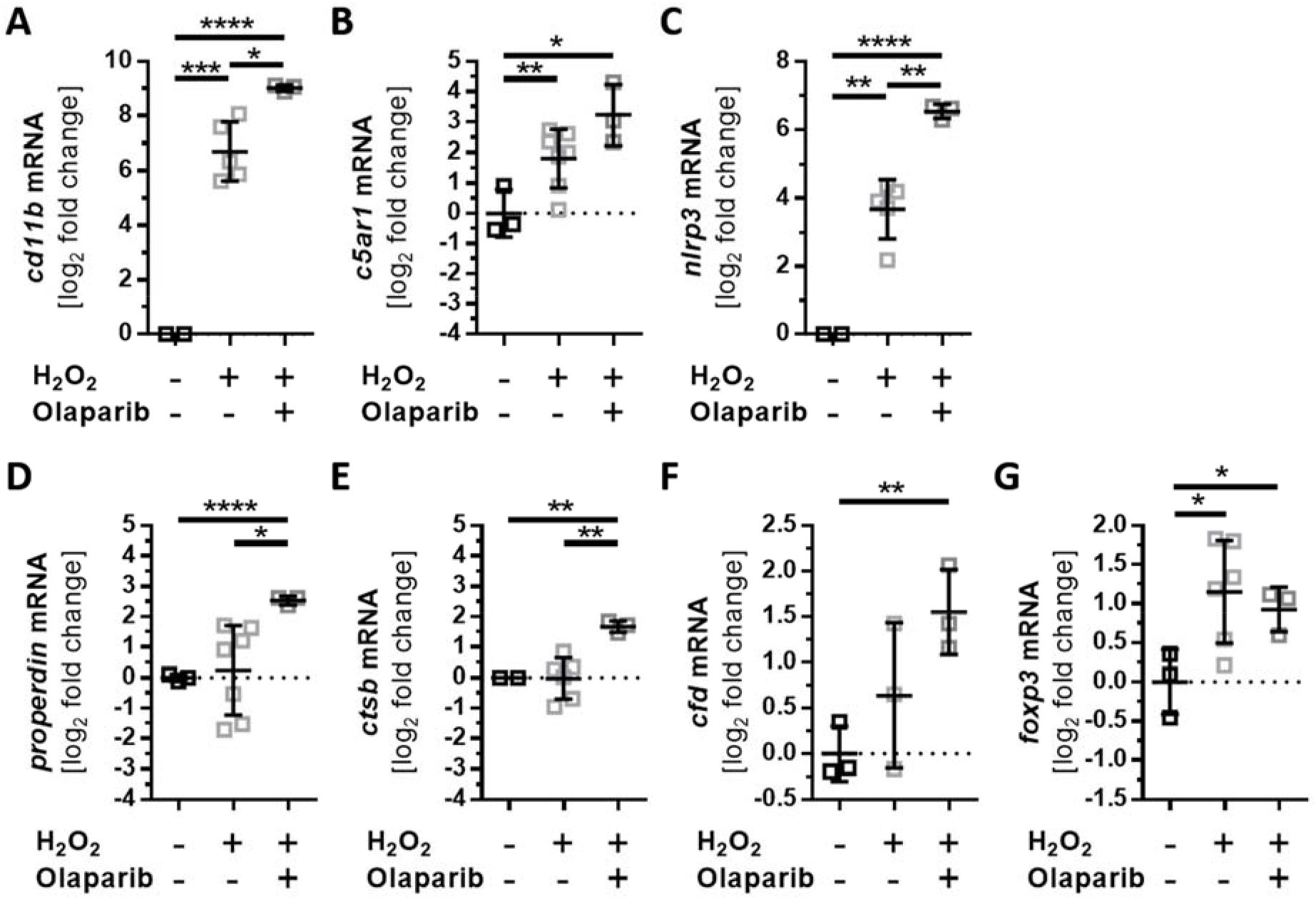
Olaparib enhanced oxidative stress dependent expression changes in ARPE-19 cells. ARPE-19 cells were treated for 4 h with H_2_O_2_ and the effect of simultaneously added olaparib on transcription was investigated. **(A)** *Cd11b*, **(C)** *nlrp3*, **(D)** *properdin* and **(E)** *ctsb* transcripts were significantly increased in olaparib-treated, stressed cells compared to stressed cells alone. **(B)** *C5aR1*, **(F)** *cfd* and **(G)** *foxp3* mRNA expression was not significantly changed in stressed ARPE-19 cells following olaparib addition. Mean with standard deviation is shown, * p≤ 0.05, ** p≤ 0.01, *** p≤ 0.001, **** p≤ 0.0001 unpaired, two-tailed, parametric t-test, dotted line depicted untreated control.

## DISCUSSION

The RPE is exposed to high-energy light and it conducts phagocytosis of oxidized photoreceptor outer segments, both is accompanied by a rapid release of reactive oxygen species [6,43,44]. Reactive oxygen species, including H_2_O_2_, are on the one hand major cellular stressors [6,45] and, on the other hand, cellular survival factors [3,46]. Antioxidants are decreased in light-exposed retinae, allowing the intra-ocular accumulation of H_2_O_2_ [47]. We used H_2_O_2_ treatment to mimic physiological oxidative stress in serum-free cultivated RPE cells to investigate the endogenous complement response in RPE cells independent from external complement sources [48,49].

Oxidative stress increased the concentration of complement regulators CFH, properdin and of the central complement protein C3 in RPE cells time-dependently without access to any extracellular complement source. Previous studies reported mostly a reduced expression of *cfh* mRNA in RPE cells exposed to oxidative stress [29–32], but these studies did not include further CFH protein analysis. Our reported CFH protein accumulation after H_2_O_2_ treatment in polarized, monolayer ARPE-19 cells, using immunohistochemistry, is in contrast to reduced CFH protein detection results in Western Blots of non-*in vivo*-like cultivated ARPE-19 cells following H_2_O_2_ treatment [31].

However, it is known that intracellular CFH can enhance the cleavage of endogenously expressed C3 by a cathepsin L (CTSL)-mediated mechanism [50]. The lysosomal protease CTSL and the central complement protein C3 concentrations were both enhanced under oxidative stress conditions in ARPE-19 cells. Previous studies of RPE cell-derived complement components only focused on *c3* mRNA expression, which was not changed under low H_2_O_2_ concentrations [51]. We went a step further and showed that C3 was retained in the RPE cells and less secreted following oxidative stress. This RPE cell dependent local accumulation of C3 was also shown for ARPE-19 cells treated with cigarette smoke [28]. If C3 is activated in the blood, this is inhibited by CFH and promoted by complement regulator properdin. We showed for the first time, that oxidative stress increased *properdin* mRNA expression in ARPE-19 cells. This resulted in a higher properdin protein concentration in these cells, which could promote cellular C3 cleavage. In summary, our data described a local production of complement proteins in RPE cells and an enhanced cellular storage of complement proteins in the cells after H_2_O_2_ treatment. This cellular accumulation suggested an autocrine, cellular function of complement proteins in RPE cells following oxidative stress.

Our studies revealed a co-localization of accumulated, endogenous C3 with complement receptor 3 (CR3, CD11b/CD18) in ARPE-19 cells exposed to oxidative stress and an increase of CR3 after 4 h. CR3 expression had been associated with inflammasome activation as a reaction to complement components or/and oxidative stress in white blood and RPE cells [52,53]. In agreement with this, the addition of H_2_O_2_to ARPE-19 cells increased a time-dependent expression of *nlrp3* and *il-1β* mRNA and subsequently enhanced the secretion of pro-inflammatory cytokines IL-1β and IL-6, which indicated an enhanced inflammasome activity. Inflammasome activation depends on reactive oxygen species and has been associated with lipid peroxidation end products and phototoxicity in RPE cells [54,55]. Involvement of the complement components in this oxidative stress response of RPE cells had been only described in relation to extracellular added anaphylatoxins so far [25], but endogenous complement of RPE cells hasn`t been suggested as potential priming factors for the inflammasome. On the one hand, we detected activated C3 cleavage products in ARPE-19 cells and previous studies showed that activated C3a can be intracellularly generated in RPE cells independent from the systemic canonical complement system [18–21]. On the other hand, C3 receptors are expressed and regulated under oxidative stress in ARPE-19 cells indicating a role of endogenous complement components in stressed ARPE-19 cells. Cellular C3 is cleaved by lysosomal CTSL [15,50] and NLRP3-inflammasome activation depended on this CTSL activity [55]. Previously, CTSL inhibition reduced inflammasome activity in ARPE-19 cells exposed to oxidative stress [56], showing the interaction of cell-specific complement component cleavage and inflammasome activity. It is already known, that endogenous C3-driven complement activation was required for the IL-1β and IL-6, as well as inflammasome activation in immune cells [57]. Our data suggest now, that this could be also a autocrine mechanism in RPE cells.

Additionally to C3, C5 has been identified as a key player in cell homeostasis [25]. The *c5aR1* receptor is expressed in RPE cells [58,59] and was increased during oxidative stress. *C5* mRNA expression was not changed, as the expression of *c3* mRNA. However, the biologically highly active C5a fragment, a ligand for C5aR1 has a half-life of approximately 1 min [60,61], due to rapid receptor binding. This rapid signalling might have interfered with our detection scheduled. C5aR1 stimulation is associated with IL-8 and VEGF-a secretion in ARPE-19 cells [58,59]. Increased secretion of these pro-angiogenic factors was also observed following the H_2_O_2_ stimuli, but the signalling pathway is not exactly known so far. In regulatory T-cells the transcription factor FOXP3 promotes the expression of IL-8 [23] and in bladder cancer cells a knock-down of *foxp3* resulted in a reduced expression of *vegf* [24]. *Foxp3* mRNA was expressed in ARPE-19 cells and increased under oxidative stress conditions. Previous studies showed, that extracellular C5a can activate FOXP3 in ARPE-19 cells, which was associated with increased IL-8 secretion [26]. We showed that this can be also due to endogenous activation of C5aR1 following oxidative stress in RPE cells.

These changes in expression and cellular complement protein accumulation following oxidative stress were time-dependent (Sup. Fig. 5). The first changes of complement receptor (CR3, C5aR1) and component (CFH, C3) levels in the RPE cells occurred after 4 h and were accompanied with changes in *nlrp3* and *foxp3* mRNA expression. Downstream alterations in properdin expression, intracellular proteases and an increase of epithelial–mesenchymal transition marker as well as loss of tight-junctions were described. This indicates that complement receptor signalling could be involved in early response of RPE cells to H_2_O_2_treatment.

Oxidative stress-related cell damage of ARPE-19 cells and retinal degeneration in mouse models for RPE degeneration as well as hereditary retinal degeneration were successfully ameliorated using olaparib in previous studies [40–42]. Olaparib is a clinically developed poly-ADP-ribose-polymerase inhibitor developed for cancer treatment by blocking the DNA-repair mechanism. ARPE-19 cells were resistant to H_2_O_2_ induced mitochondrial dysfunction and to energy failure, when olaparib was added [40]. We ask the question if olaparib can also normalize complement-associated pro-inflammatory expression profiles in H_2_O_2_-treated cells. Surprisingly, olaparib accelerated the effect of oxidative stress in RPE cells and enhanced the expression of complement receptors, complement components and the *nlrp3* mRNA. This shows that endogenous complement-related, pro-inflammatory response of ARPE-19 cells could be correlated with defective DNA repair mechanisms.

## CONCLUSION

Oxidative stress and activation of the complement system cause retinal degeneration, but the mechanism behind this is still a matter of investigation. We showed for the first time, that oxidative stress can increase endogenous RPE cell complement components and receptors and that the process was associated with release of pro-inflammatory and pro-angiogenic factors. Our data offer a stepping stone for numerous further investigations regarding the function of a cell-associated complement system in the RPE. Many questions were raised during this project: How are the complement components activated? What is (are) the signalling pathway(s) of the complement receptors independent from external complement sources? How are inflammasome regulation and FOXP3 activity modulated by endogenous complement components in RPE cells? Can endogenous complement factors be targeted to affect cell-associated signalling pathways? These new perspectives will hopefully help to decipher the function of intracellular complement components in retinal health and disease and offer new strategies for treatment of retinal degeneration.

## METHODS

### Cell culture and treatment

Human ARPE-19 cells (passage 39; American Type Culture Collection) were cultivated for 6 days in cell culture flasks with DMEM/F12 (Sigma-Aldrich) and 10% fetal calf serum (FCS; PanBiotech) and 1% penicillin/ streptomycin (37°C, 5% CO_2_). Cells were trypsinized (0.05% trypsin/ 0.02% EDTA) and seeded in a concentration of 1.6 × 10^5^ cells/cm^2^ (passage 39) on mouse laminin (5 µg/cm^2^, Sigma-Aldrich) coated 0.4 μm pore polyester membrane inserts (Corning). Cells were cultivated for 4 weeks with apical and basal media exchanges (first day medium with 10% FCS, remaining time medium with 5% FCS were used). Before treatment FCS concentration was reduced within 3 days from 5% to 0%. ARPE-19 cells were treated either with 0.5 mM H_2_O_2_ for 1, 4, 24 and 48 h, or 0.5 mM H_2_O_2_ and 0.01 mM Olaparib (Biomol, Hamburg, DE) for 4 h.

### Immunohistochemistry and TUNEL assay

PBS (Sigma-Aldrich) washed, paraformaldehyde (4%, 20 min; Merck) fixated ARPE-19 cells were permeabilized (PBS/ 0.2% Tween20 (PBS-T), 45 min) and unspecific bindings were blocked (3% BSA (Carl Roth)/PBS-T, 1 h). Antigens were detected using primary antibody (Sup. Table 1, overnight, 3% BSA/PBS-T) and fluorescence-conjugated anti-species antibody (Sup. Table 1, 45 min, 3% BSA/PBS). HOECHST 33342 (1:1000) stained DNA. Cells were covered with fluorescenting mounting medium (Dako, Agilent). Images were taken with a confocal microscope (Zeiss).

The TUNEL assay was performed with DeadEnd™ Fluorometric TUNEL System (Promega) on paraformaldehyde fixated, washed and permeabilized (0.2% Triton X-100 in PBS) cells. Images were taken with confocal microscope a by Zeiss.

### RT-qPCR

mRNA was isolated using the NucleoSpin^®^ RNA/Protein kit (Macherey-Nagel). Purified mRNA was transcribed into cDNA with the QuantiTect^®^Reverse Transcription Kit (Qiagen). Transcripts of complement components, receptors and inflammation-associated markers were analyzed using the Rotor-Gene SYBR^®^Green PCR Kit either with QuantiTect Primer Assays (Sup. Table 2), or in-house designed primer pairs (Metabion) described in Sup. Table 3 in the Rotor Gene Q 2plex cycler (Qiagen).

### Western Blot

Proteins were purified using RIPA buffer (Sigma-Aldrich) with protease and phosphatase inhibitors (1:100, Sigma-Aldrich). Samples were dissolved in reducing Laemmli sample buffer and denatured (95 °C, 10 min). Samples were separated in a 12% SDS-PAGE and transferred on to an activated polyvinylidene difluoride membrane using a wet blotting system. Membranes were blocked (1 h, 5% BSA/PBS-T) and incubated with the primary antibody (Sup. Table 1, overnight, 5% BSA/PBS-T). Peroxdiase-conjugated anti-species antibodies were used for detection (Sup. Table 1, 1 h, PBS-T). WesternSure PREMIUM Chemiluminescent Substrate (LI-COR) visualized the antigen in the Alpha Innotech Fluor Chem FC2 Imaging System.

### Multiplex-Immunoassays

Cytokine concentration of basal and apical supernatants of treated and untreated ARPE-19 cells were determined according to the protocol of a custom ProcartaPlex^®^ multiplex immunoassay kit (ThermoFisher). Complement components in the cellular supernatant were quantified using the MILLIPLEX MAP Human Complement Panel (Merck). The read out of the multiplex assay was performed in a Magpix instrument (Luminex).

### Statistics

Statistical analysis was performed using GraphPad Prism 7 (GraphPad Software Inc.).

## DATA AVAILABILITY

Original data supporting the findings of this study are available from the corresponding author upon reasonable request.

## ACKNOWLEDGEMENTS

This project was supported by the Velux Foundation (Proj. Nr. 1103) to DP and VE. We thank Renate Foeckler, Andrea Dannullis and Elfriede Eckert for excellent technical support.

## AUTHOR CONTRIBUTIONS

TT, NS, VE and DP designed research; TT, NS, MR, KK, VE and DP performed research; TT, NS, MR, KK, VE and DP analysed and interpreted the data; TT, NS, VE and DP wrote the manuscript. All authors provided input to edit the manuscript.

## COMPETING INTERESTS

The authors declare no competing interests.

## SUPPLEMENT

**Sup. Fig. 1.**
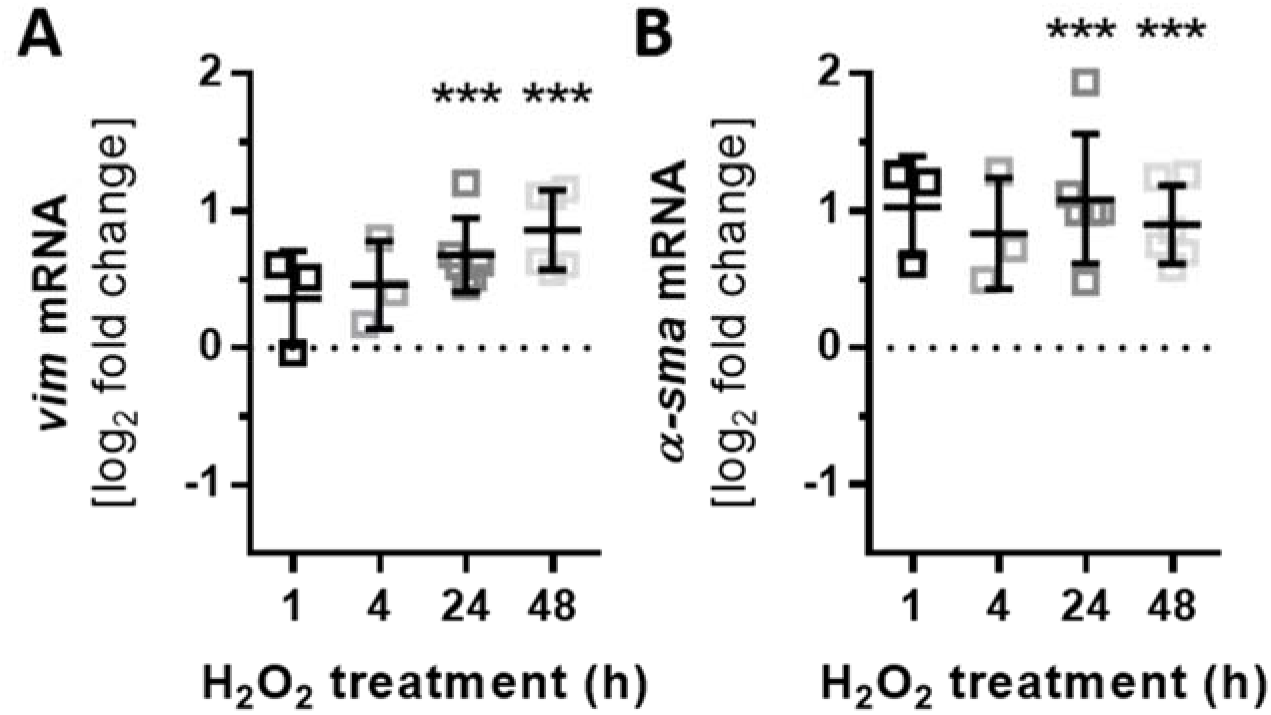
H_2_O_2_ treatment increased expression of epithelial-mesenchymal transition markers in ARPE-19 cells. ARPE-19 cells were treated either for 1, 4, 24 or 48 h with H_2_O_2_. **(A)** *vim* and **(B)** *α-sma* transcription was significantly increased after 24 h compared to the untreated control. Mean with standard deviation is shown, *** p< 0.001, unpaired, two-tailed, parametric t-test, dotted line depicts untreated control.

**Sup. Fig. 2.**
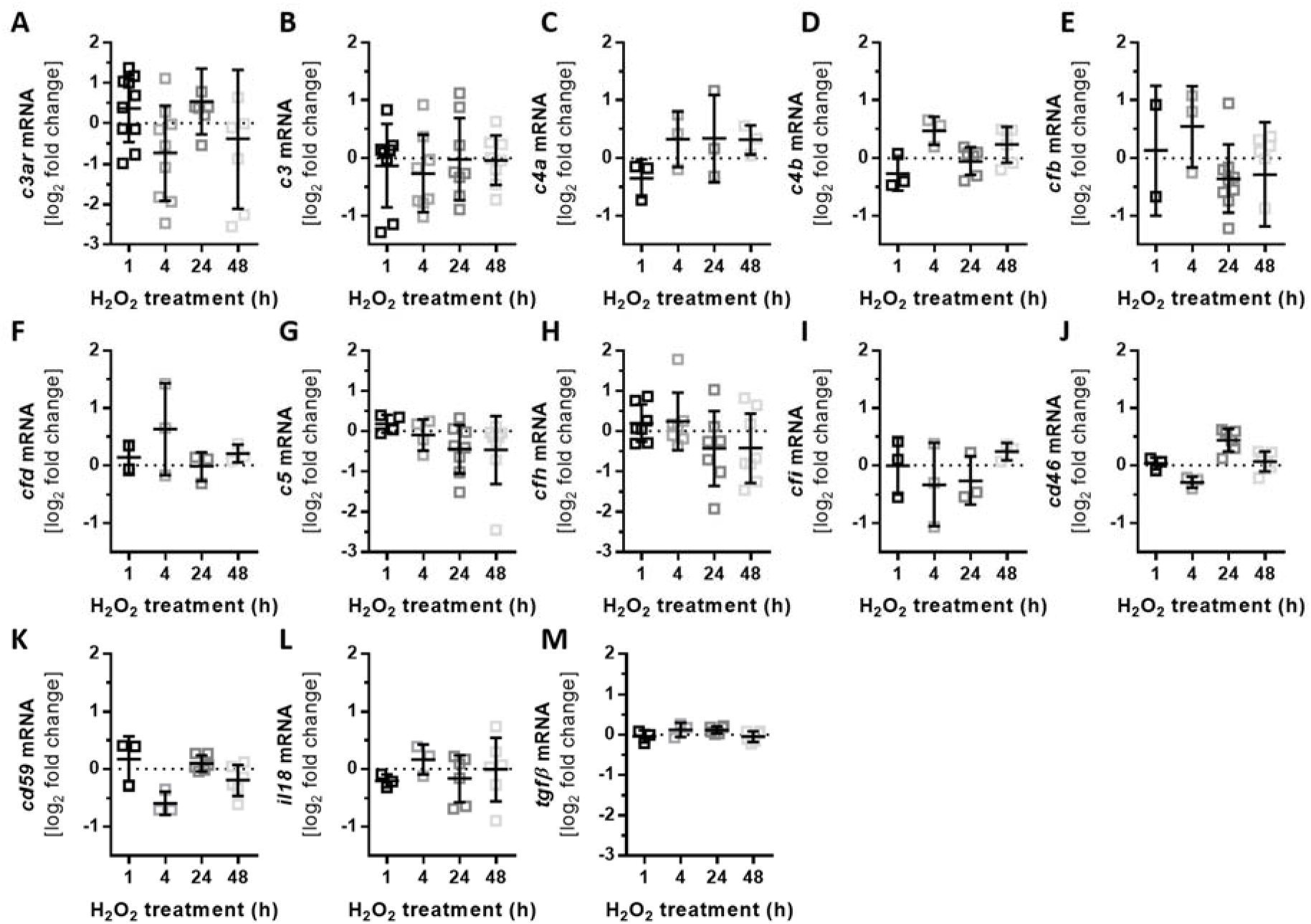
H_2_O_2_ treatment did not influence the transcription levels of several genes in ARPE-19 cells. ARPE-19 cells were treated either for 1, 4, 24 or 48 h with H_2_O_2_. mRNA levels were not significantly changed for: **(A)** *c3ar*, **(B)** *c3*, **(C)** *c4a*, **(D)** *c4b*, **(E)** *cfb*, **(F)** *cfd*, **(G)** *c5*, **(H)** *cfh*, **(I)** *cfi*, **(J)** *cd46*, **(K)** *cd59*, **(L)** *il18* and **(M)** *tgfβ*. Mean with standard deviation, dotted line untreated control.

**Sup. Fig. 3.**
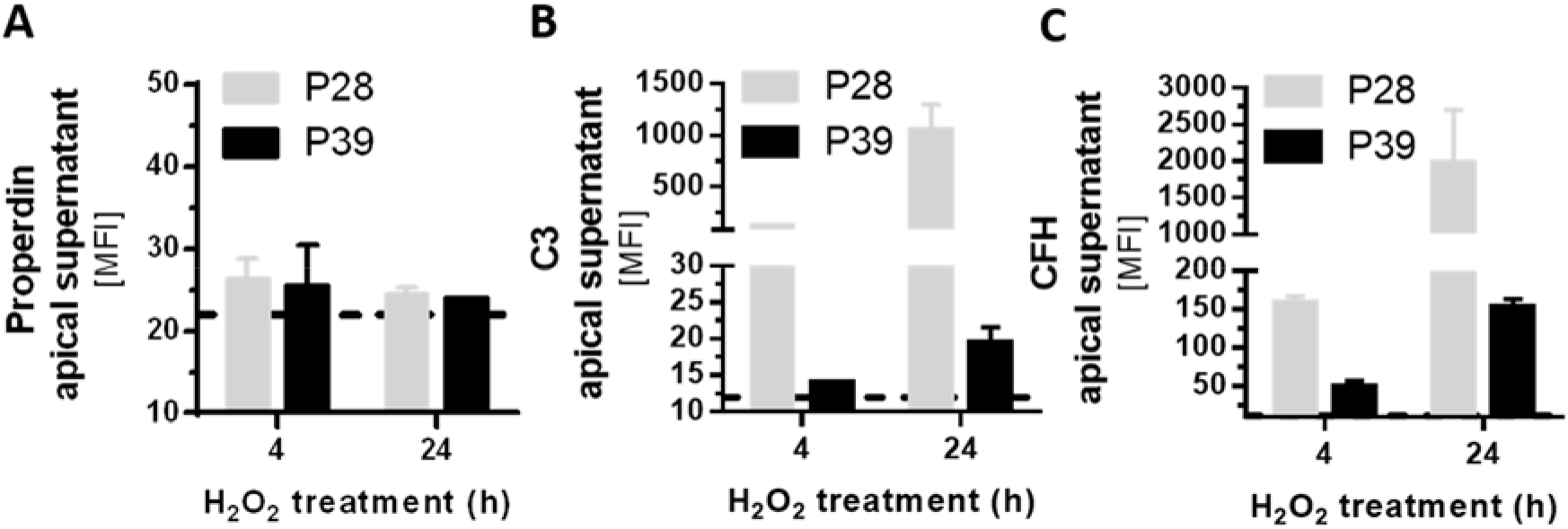
Increased complement component secretion in lower passage number of ARPE-19 cells compared to higher passage number. ARPE-19 cells with passages 28 or 39 (latter used in the rest of this study) were treated either for 4 or 24 h with H_2_O_2_. The protein concentration of **(A)** properdin, **(B)** C3 and **(C)** CFH was determined in the apical supernatant using a multiplex immunoassay. **(A)** Properdin was not secreted by ARPE-19 cells of varied passages. ARPE-19 cells with lower passage number secreted more **(B)** C3 and **(C)** CFH than ARPE-19 cell with higher passage number into the apical supernatant.

**Sup. Fig. 4.**
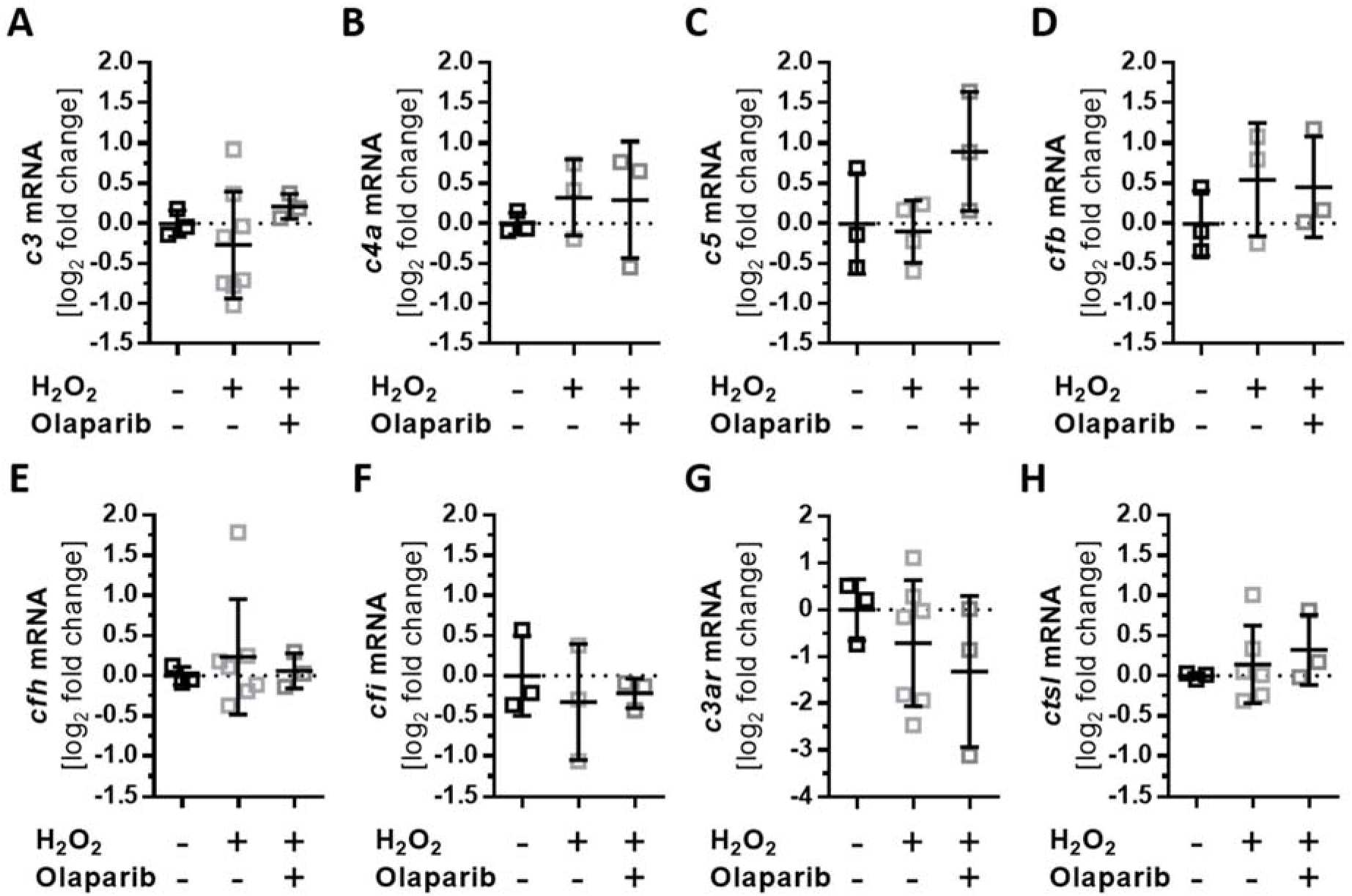
Stable expression of complement components and related genes after Olaparib and oxidative stress treatment in ARPE-19 cells. ARPE-19 cells were treated for 4 h with H_2_O_2_ and the effect of simultaneously added olaparib on transcription was investigated. **(A)** *c3*, **(B)** *c4a*, **(C)** *c5*, **(D)** *cfb*, **(E)** *cfh*, **(F)** *cfi*, **(G)** *c3aR* and **(H)** *ctsl* did not significantly change in stressed ARPE-19 cells following olaparib addition.

**Sup. Fig. 5.**
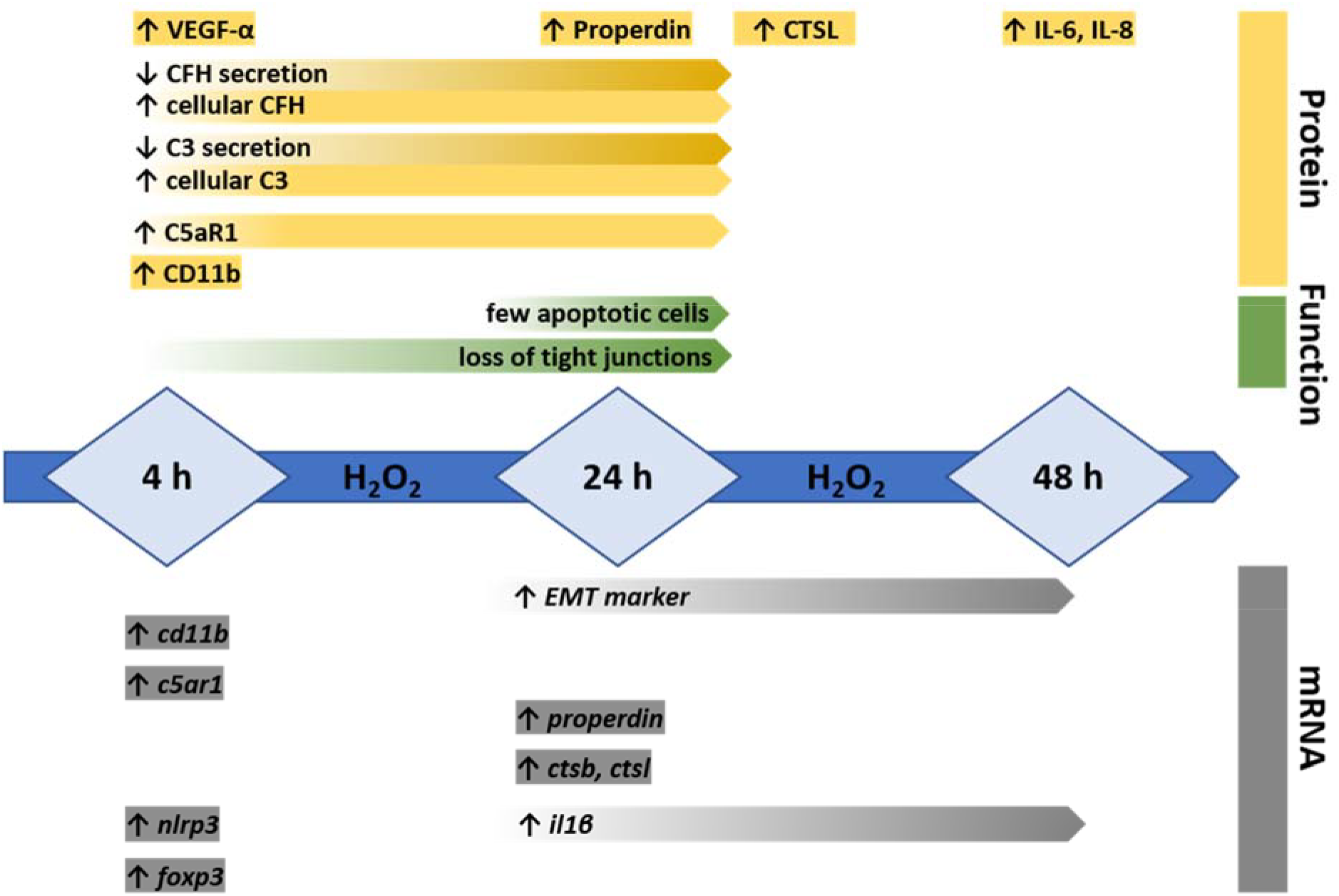
Time-dependent changes of H_2_O_2_ treatment in ARPE-19 cells. ARPE-19 cells were treated for 4, 24 and 48 h with H_2_O_2_. Changes in mRNA expression (grey), function (green) and on protein level (yellow), which are described in this manuscript, are summarized in this scheme.

**Sup. Table 1:**
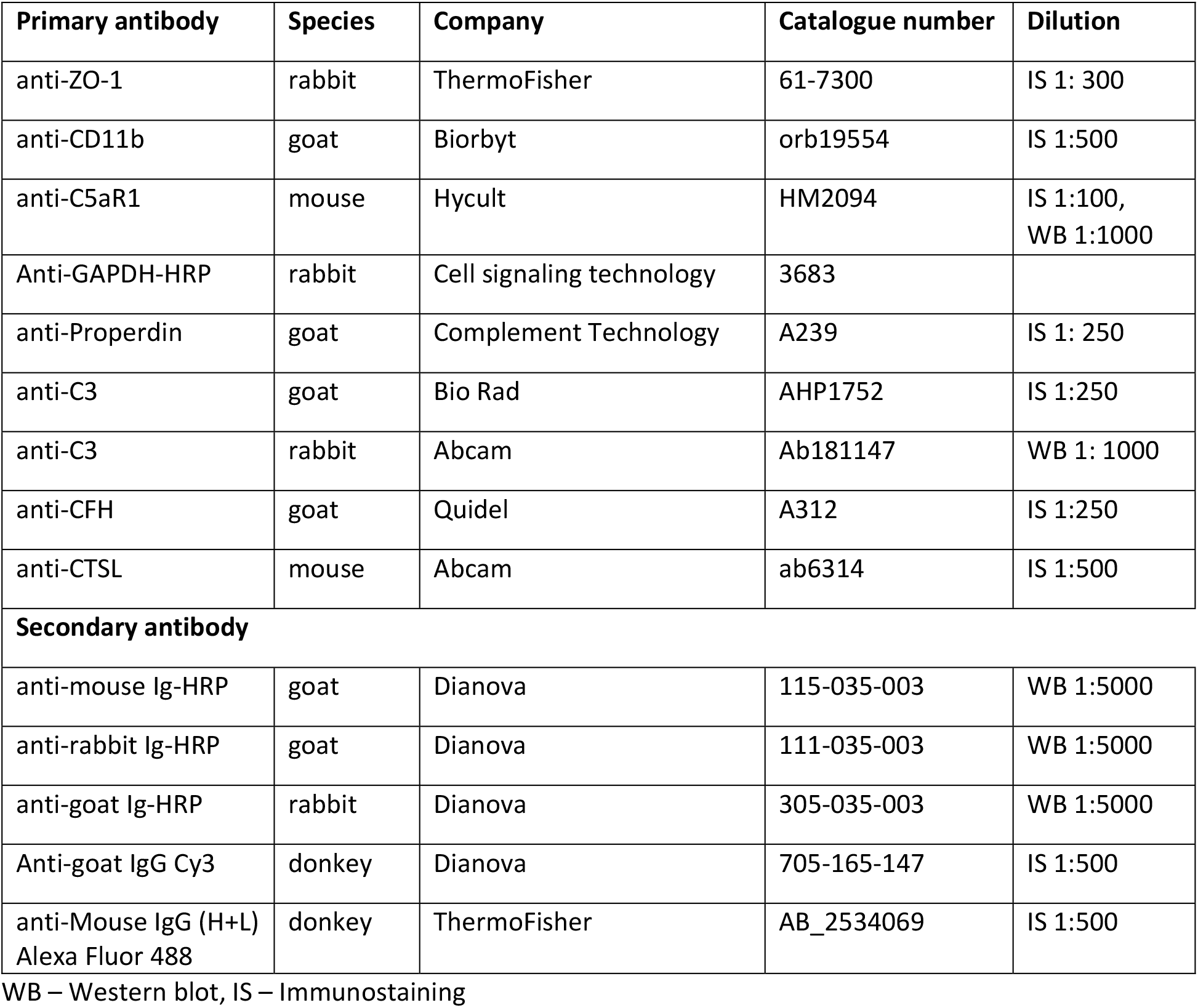
Primary and secondary antibodies

**Sup. Table 2:** QuantiTec PrimerAssays

| mRNA transcript | name | Catalogue number |
| --- | --- | --- |
| *gapdh* | Hs_GAPDH_1_SG | QT00079247 |
| *c3* | Hs_C3_1_SG | QT00089698 |
| *c3ar* | Hs_C3AR1_1_SG | QT00090398 |
| *cd11b* | Hs_ITGAM_1_SG | QT00031500 |
| *c4a* | Hs_C4A_1_SG | QT00237160 |
| *c4b* | Hs_C4B_1_SG | QT00237167 |
| *c5* | Hs_C5_1_SG | QT00088011 |
| *c5ar1* | Hs_C5R1_1_SG | QT00997766 |
| *cd46* | Hs_MCP_1_SG | QT00073689 |
| *cd59* | Hs_CD59_1_SG | QT00035952 |
| *cathepsin b* | Hs_CTSB_1_SG | QT00088641 |
| *cathepsin l* | Hs_CTSL_1_SG | QT01664978 |
| *complement factor b* | Hs_BF_1_SG | QT00012138 |
| *complement factor d* | Hs_CFD_1_SG | QT00212191 |
| *complement Factor h* | Hs_CFH_1_SG | QT00001624 |
| *complement Factor i* | Hs_CFI_1_SG | QT00213794 |
| *complement Factor p* | Hs_CFP_1_SG | QT00010514 |
| *nlrp3* | Hs_NLRP3_1_SG | QT00029771 |
| *forkhead-box-protein P3* | Hs_FOXP3_1_SG | QT00048286 |

**Sup. Table 3:**
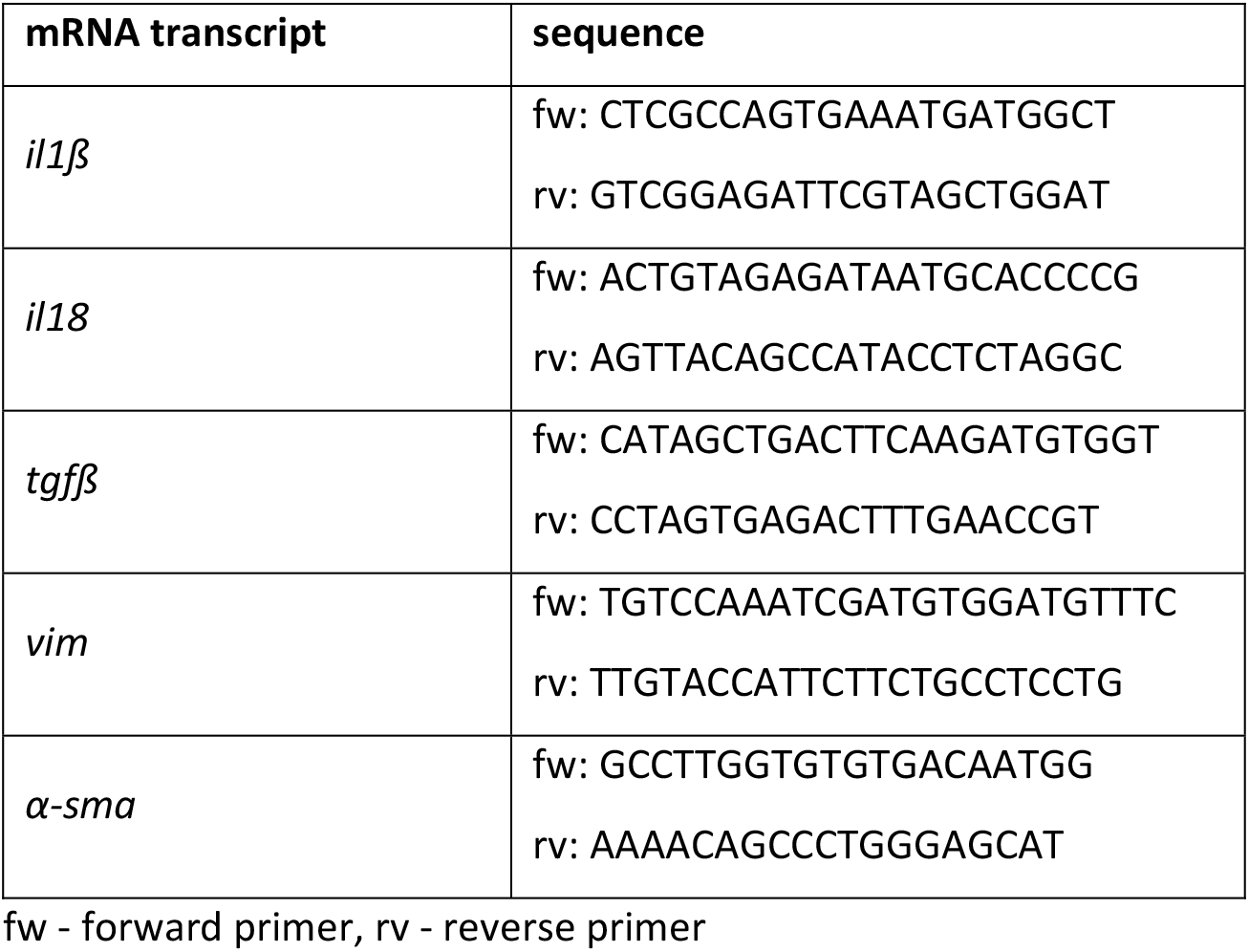
In-house designed RT-qPCR primers

